# Guided self-organization recapitulates tissue architecture in a bioengineered brain organoid model

**DOI:** 10.1101/049346

**Authors:** Madeline A. Lancaster, Nina S. Corsini, Thomas R. Burkard, Juergen A. Knoblich

**Affiliations:** IMBA - Institute of Molecular Biotechnology of the Austrian Academy of Science Vienna 1030, Austria; MRC Laboratory of Molecular Biology, Cambridge Biomedical Campus, Cambridge, UK

**Keywords:** Neurogenesis, human brain development, organoid, scaffold, patterning, neuronal migration, fetal alcohol syndrome

## Abstract

Recently emerging methodology for generating human tissues *in vitro* has the potential to revolutionize drug discovery and disease research. Currently, three-dimensional cell culture models either rely on the pronounced ability of mammalian cells to self organize *in vitro*^1-6^, or use bioengineered constructs to arrange cells in an organ-like configuration^7,8^. While self-organizing organoids can recapitulate developmental events at a remarkable level of detail, bioengineered constructs excel at reproducibly generating tissue of a desired architecture. Here, we combine these two approaches to reproducibly generate micropatterned human forebrain tissue while maintaining its self-organizing capacity. We utilize poly(lactide-co-glycolide) copolymer (PLGA) fiber microfilaments as a scaffold to generate elongated embryoid bodies and demonstrate that this influences tissue identity. Micropatterned engineered cerebral organoids (enCORs) display enhanced neuroectoderm formation and improved cortical development. Furthermore, we reconstitute the basement membrane at later stages leading to characteristic cortical tissue architecture including formation of a polarized cortical plate and radial units. enCORs provide the first *in vitro* system for modelling the distinctive radial organization of the cerebral cortex and allow for the study of neuronal migration. We demonstrate their utility by modelling teratogenic effects of ethanol and show that defects in leading process formation may be responsible for the neuronal migration deficits in fetal alcohol syndrome. Our data demonstrate that combining 3D cell culture with bioengineering can significantly enhance tissue identity and architecture, and establish organoid models for teratogenic compounds.

---

Due to the nature of their self-organization, the current usability of organoids is limited by high variability, random tissue identity and incomplete morphological differentiation^9,10^. To identify sources of variability in brain organoids^11^, we examined five independent batches of cerebral organoids at various time points. We found that batch-to-batch variation started during neural induction, when the efficiency of polarized neural ectoderm formation varied between 30% and 100% of organoids per batch (Supplemental Fig. 1). Organoids that conformed well with the bright field microscopy criteria laid out in our protocol^11^ contained well-organized, apicobasally polarized neural epithelium around the surface of the organoid with only sparse cells staining positive for other germ layers (Supplemental Fig. 2a). Organoids classified as “suboptimal”, however, displayed less well-organized architecture with a greater extent of endodermal and mesodermal identities (Supplemental Fig. 2a). During later stages, these organoids contained tissues with non-neural morphologies such as putative cartilage, mesenchyme and squamous epithelia (Supplemental Fig. 2b). Immunostaining identified occasional definitive endoderm and other non-neural E-cadherin positive regions (Supplemental Fig. 2c), suggesting that inconsistent neural induction efficiency might be a main source of variability.

Blocking both BMP and Activin/Nodal signalling^12-14^ as well as addition of retinoids^15^ have been shown to efficiently induce neural identity. Indeed, organoids treated with retinoids and the SMAD inhibitors dorsomorphin and SB-431542 displayed increased neuroepithelial tissue during neural induction (Supplemental Fig. 3a). During later stages, however, drug addition resulted in small rosette-like structures that prematurely displayed neural processes upon embedding in matrigel (Supplemental Fig. 3b) rather than the neural tube-like buds typical of cerebral organoids. Thus, acceleration of neural induction *in vitro*^13^ compromises 3D tissue architecture possibly due to an effective omission of early embryonic events such as ectoderm to neural ectoderm transition.

While gastrulating human embryos are disc-shaped^16^, most brain organoid protocols start from spherical EBs. These are generated by pipetting hPSCs into non-adhesive plastic microplates and aggregating the cells at the bottom of U-shaped wells^17-19^. Low surface to volume ratio in the resulting spherical EBs may limit the extent of neuroectoderm formation, which develops on the outside of the EB. It is known that micropatterned substrates can influence pluripotency^20^ and germ layer identity during PSC differentiation^21^. However, this approach is incompatible with current organoid protocols that all require non-adhesion to a surface and extensive cell-cell contact for self-organization. We therefore used a floating scaffold to pattern the organoids at the EB stage from the inside. Furthermore, in order to limit cell contact to the scaffold and thus maintain dense cell-cell contact, we used micrometer-scale individual filaments that would contact only the innermost layer of cells of the EB. Filaments were obtained by mechanical dispersion from braided fibers of PLGA 10:90 (Supplemental Fig. 4a-c), a bio-compatible material that is absorbed in living tissue by hydrolysis within 8-10 weeks^22^. 5-10 microfilaments were collected in a random configuration at the bottom of a low-attachment round-bottom microwell and seeded with 18,000 hPSCs (Supplemental Fig. 4d), a ratio that results in only 5-10% of cells in direct contact with the filament.

The resulting micropatterned engineered cerebral organoids (enCORs) displayed elongated morphologies but maintained typical dense cell composition. Like spherical EBs, they progressively grew in size and exhibited clearing along the edges with eventual formation of polarized neural ectoderm. However, the neural ectoderm was dramatically elongated (Figure 1b) and the efficiency of neuroectoderm formation was much improved with consistent neural induction in all of five independent preparations examined (Supplemental Fig. 4e). Staining of early stage micropatterned EBs for germ layer markers revealed consistent generation of polarized neuroepithelium with concomitantly decreased amounts of endoderm and mesoderm identities (Figure 1c and Supplemental Fig. 5a). In addition, expression analysis for pluripotency and germ layer markers by RT-PCR, revealed a decrease in non-neural identities (Supplemental Fig. 5b, c). Finally, enCORs displayed fewer morphological features of other germ layers such as the formation of early fluid-filled cysts (Supplemental Fig. 5d).

**Figure 1.**
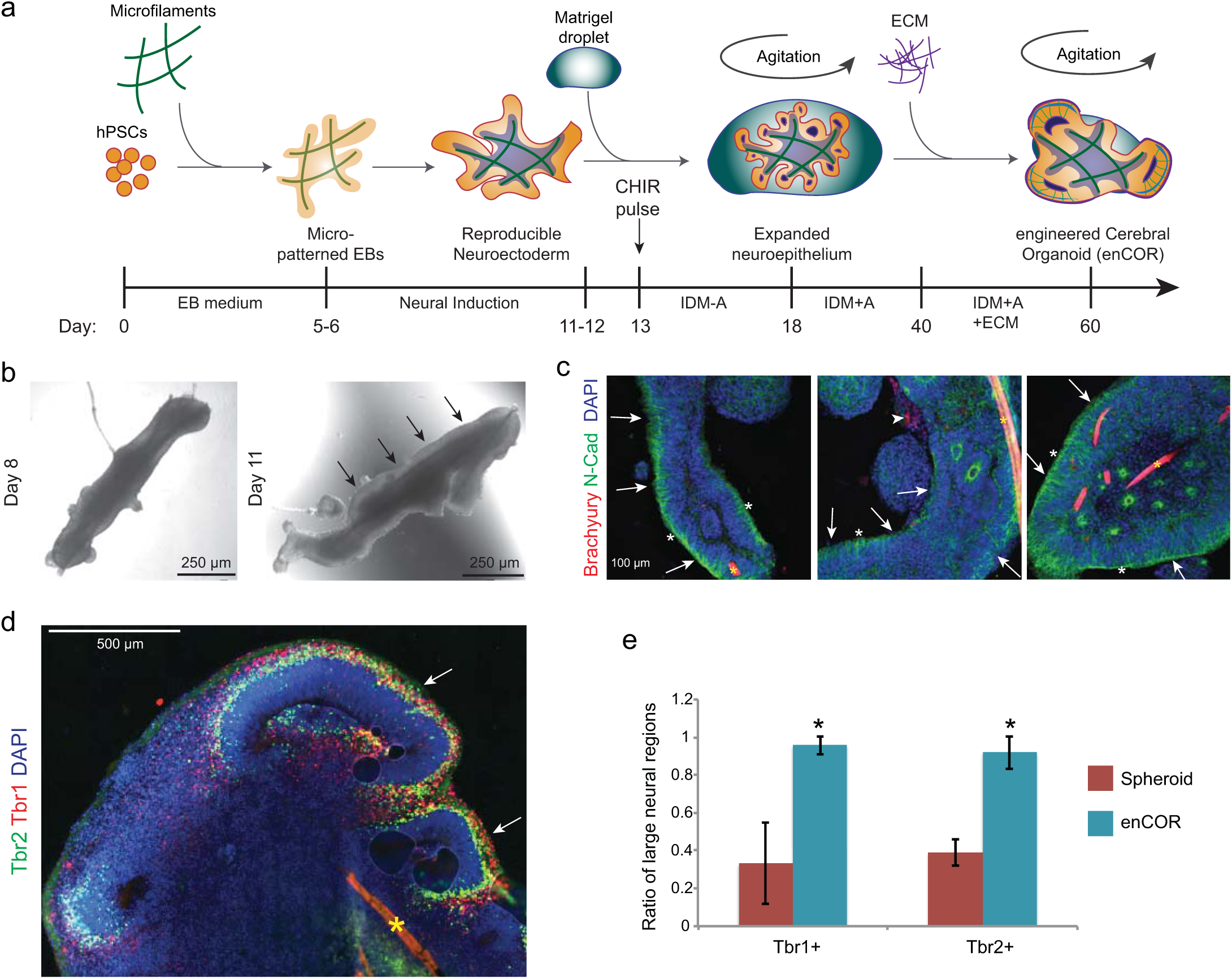
Engineered cerebral organoids reproducibly generate organized neural tissue. **a**. Schematic of the method for generating engineered cerebral organoids (enCORs). Timeline and media used (methods) are shown at the bottom. **b**. Micropatterned EBs at two time points during neural induction, showing clearing along the edges and polarized neural ectoderm (arrows). **c**. Immunohistochemical staining of day 10 micropatterned EBs for the germ layer markers Brachyury for mesoderm and N-Cadherin for neural ectoderm. Note the prevalence of polarized neural epithelia (arrows) displaying the apical domain on the surface (white asterix) with only occasional mesoderm identities (arrowhead). The microfilament can be seen as an autofluorescent rod (yellow asterix). **d**. Staining for dorsal cortical markers Tbr2 and Tbr1 identifies large lobes of brain tissue within microfilament patterned organoids treated with CHIR to be dorsal cortex (arrows). **e**. Quantification of the ratio of lobes of brain tissue that were positive for Tbr1 or Tbr2 in spheroid or microfilament patterned organoids. \**P*<0.05, Student’s *t*-test, n=8 organoids from 3 batches for spheroid, n=17 organoids from 6 batches for ENCOR.

Since the neuroepithelial buds that formed upon Matrigel embedding were less continuous in enCORs (Supplemental Fig. 6a), we added a short 3-day pulse of the GSK3beta inhibitor CHIR99021 (CHIR) after neuroepithelial budding. Wnt pathway activation induces lateral expansion of cortical neuroepithelium^23^ and indeed, addition of CHIR resulted in larger lumens with surrounding continuous neuroepithelium (Supplemental Fig. 6b). These features were maintained during later stages, when enCORs contained large lobes of brain tissue (Supplemental Fig. 6c) but only few regions expressing markers for endoderm and non-neural epithelia (Supplemental Fig. 6d), in contrast to spherical organoids (Supplemental Fig. 2c). In addition, because Wnt is a dorsalizing factor in the CNS^24^, CHIR addition during lateral expansion also influenced regional identity. enCORs contained large Emx1+ regions (Supplemental Fig. 6e), consistent with dorsal cortex. Virtually all large regions stained positive for dorsal cortical markers Tbr1 and Tbr2 (Figure 1d, e, Supplemental Fig. 6f), while only approximately 30% of the large brain lobes expressed these markers in sub-optimal spherical organoids. Thus, late CHIR treatment not only expands the neuroepithelium, but also promotes reliable dorsal forebrain regionalization.

Improved brain patterning was confirmed by staining for various other brain region identities. Staining for the broad forebrain marker Foxg1 and Pax6 (Supplemental Fig. 7a) revealed the presence of forebrain regions in spherical organoids that were not Pax6 positive, whereas enCORs displayed costaining, consistent with a dorsal cortical identity. Staining for the ventral forebrain marker Gsh2 and the more specific dorsal marker Tbr2 (Supplemental Fig. 7b) revealed the presence of large ventral regions in the spheroids, whereas enCORs displayed large dorsal cortical regions, and only occasional smaller ventral regions. Besides the cortex, other dorsal forebrain derived regions such as choroid plexus and hippocampus where also evident in enCORs (Supplemental Fig. 7c, d). In constrast, spherical organoids also contained more caudal regions like spinal cord marked by Pax2 (Supplemental Fig. 7e). Thus, together with late GSK3beta inhibition, micropatterning of cerebral organoids results in homogeneous dorsal forebrain tissue with little contamination from other germ layers and brain regions.

To further examine the effect of micropatterning and CHIR addition we analyzed gene expression at 20 and 60 days in three enCORs and three spherical organoids (Supplementary Table 1). 20 day enCORs were enriched for GO terms “neurological system” and “multicellular organismal processes”, while other organ development like digestive tract, muscle, skeletal system and mesoderm were decreased (Figure 2a). At 60 days, we observed GO term enrichment for nervous system development and transcription while digestive tract, heart, muscle skeletal system, and synaptic transmission were decreased. The decrease in synaptic genes at 60 days suggests a delay in neuronal maturation perhaps due to extended progenitor expansion upon CHIR addition. Hierarchical cluster analysis revealed several gene clusters displaying specific patterns of differential expression (Figure 2b, Supplemental Figure 8, Supplementary Table 2). Cluster 1 was upregulated in 60 day enCORs and enriched for forebrain and cortical differentiation. Clusters 4 and 5 were increased or unchanged at 20 but decreased at 60 days and were enriched for nervous system development and synaptic transmission. Finally, Clusters 6 and 7 were decreased at 20 days and also decreased or unchanged at 60 days. They were enriched for more caudal expression such as spinal cord and hindbrain, consistent with the effect of micropatterning and CHIR addition on forebrain patterning.

**Figure 2.**
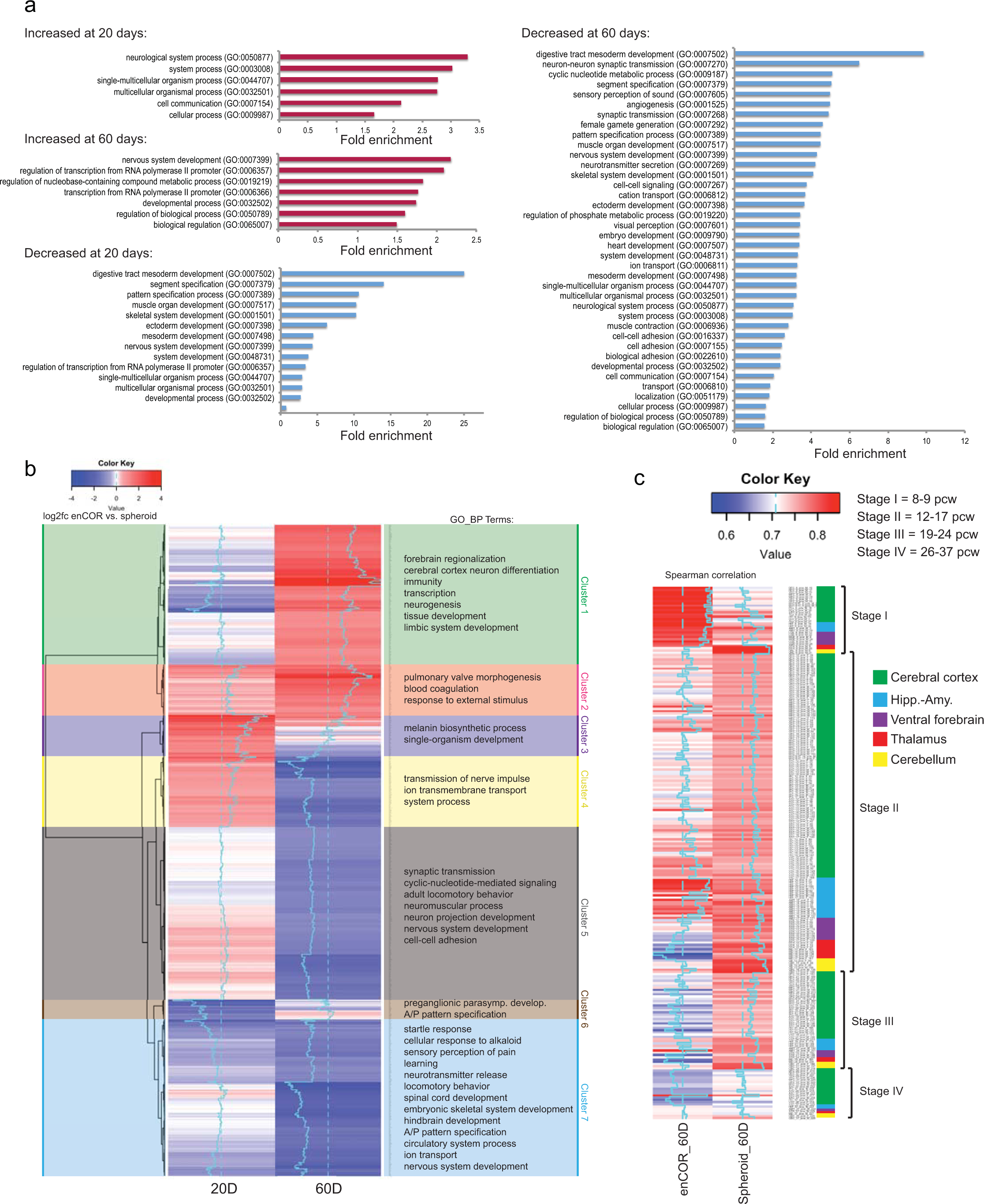
enCORs generate pure neural tissue and are preferentially dorsal forebrain. **a**. Fold enrichment of GO Slim Biological Process terms for genes increased or decreased in micropatterned organoids at 20 and 60 days. **b**. Heatmap of hierarchical clustering of genes according to log2fc values at 20 days and 60 days in micropatterned versus spheroid organoids. Clusters are marked and GO biological process term summaries of terms identified (Supplemental Fig. 8) shown at right. **c**. Heatmap of Spearman correlation coefficients between micropatterned or spheroid 60 day samples and all brain regions at fetal timepoints from the Allen brainspan transcriptome, sorted by anterior-posterior regional identity and four stages of development.

We next assessed expression of specific germ layer or brain patterning markers (Supplemental Fig. 9). The pluripotency markers Oct4, Klf4, and Nanog were decreased in enCORs. Neuroectodermal markers appeared unchanged whereas mesendodermal markers such as Sox17, T, Mixl1, and Foxa2 were decreased. Furthermore, the rostral brain marker Foxg1 was dramatically increased, whereas caudal markers such as En2, Gbx2 and Hox genes were decreased. Finally, dorsal forebrain markers such as Emx1, Tbr1 and Tbr2 were increased, while ventral markers showed low expression in both. These findings suggest a more rostral-dorsal brain identity in enCORs.

Finally, we compared genes differentially expressed between 60 day spheroid organoids and enCORs to gene expression in the human developing brain using Allen BrainSpan Atlas^25^ (Figure 2c). enCORs most closely matched the dorsal forebrain identities of the human brain at early gestation, specifically 8-9 weeks post-conception. Spherical organoids instead showed the highest correlation with more caudal regions, specifically the thalamus and cerebellum, and showed a broader correlation with later time points. These findings are consistent with the effect of micropatterning and CHIR addition on dorsal forebrain regional identity and delayed maturation.

Neurons in the developing cortex show radially aligned morphology and form a dense band called the cortical plate. This is thought to reflect their organization into radial units^26^, a prerequisite for the formation of functional neuronal columns in the adult cortex ^27,28^. So far, cortical plate formation has not been successfully modelled *in vitro* where neurons are instead randomly oriented once outside the progenitor zones. This might be due to the absence of a basement membrane, which is thought to be generated by the overlying non-neural mesenchyme^29,30^ not present in organoids. Indeed, staining for laminin revealed a well-formed basement membrane in early organoids before neurogenesis, but later punctate staining, suggesting breakdown of the membrane upon generation and basal migration of neurons (Figure 3a). We therefore reconstituted the basement membrane by including exogenous extracellular matrix (ECM) components in the form of dissolved Matrigel in the media. Indeed, this maintained a thick laminin rich basement membrane that was notably outside the migrating neurons (Figure 3b), suggesting that it was not broken down by their generation and basal migration. Bright field imaging revealed a band of density in cortical regions that was absent in organoids lacking dissolved ECM (Supplemental Fig. 10a). Subsequent sectioning and histological staining revealed a radialized basal layer consistent with cortical plate morphology (Figure 3c). Indeed, immunohistochemical staining revealed it was positive for the neural markers Ctip2, and Map2, with a band of lower Map2 staining in cell bodies that is typical of the cortical plate *in vivo* (Figure 3d and Supplemental Fig. 10b).

**Figure 3.**
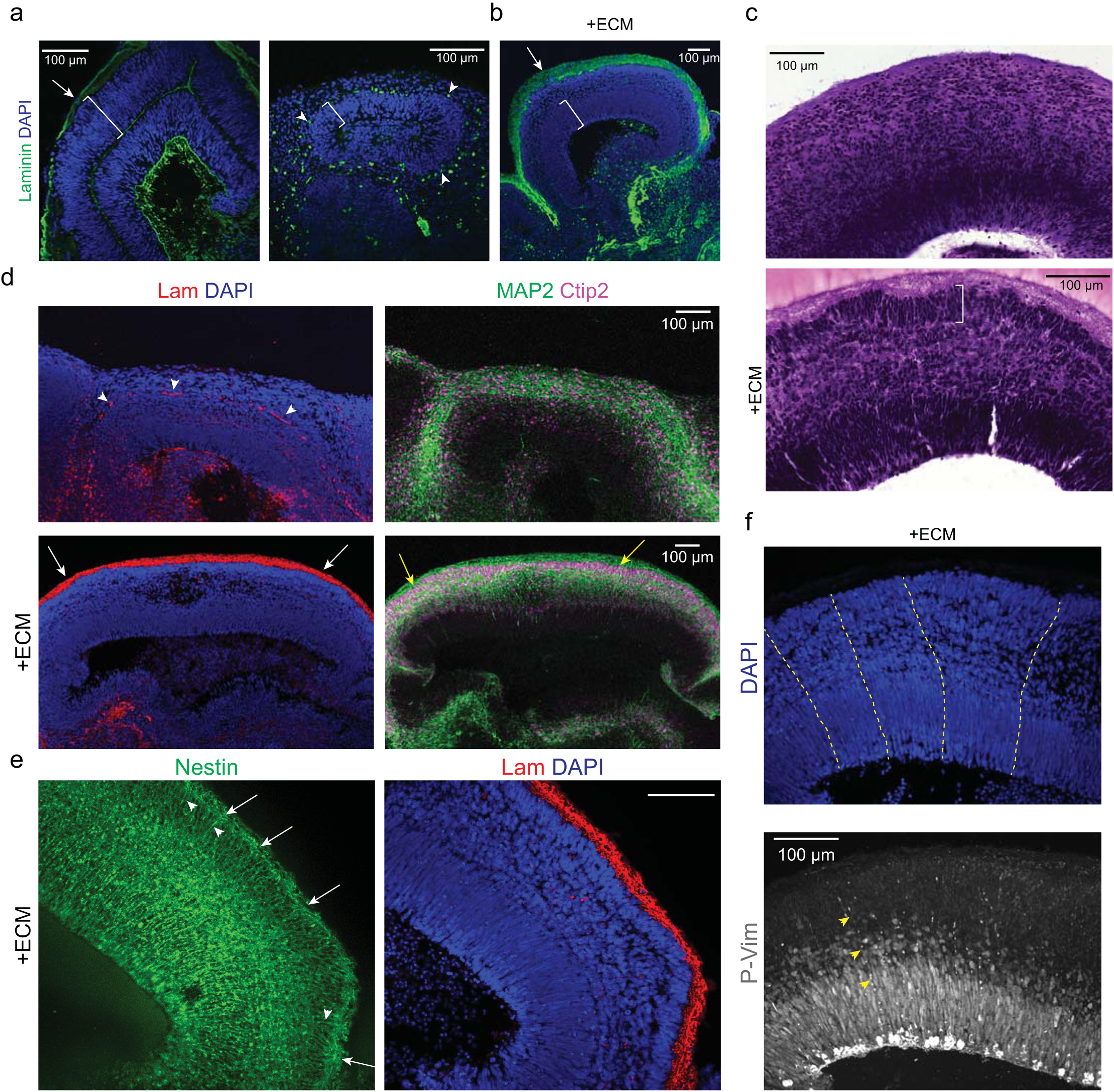
enCORs with reconstituted basement membrane form cortical plate and radial units. **a**. Staining for the basement membrane component laminin (green) in spheroid organoids. Note the presence of the basement membrane surrounding the early neuroepithelium before the generation of neurons (first panel, arrow), whereas upon neuron generation only sparse labeling remains (arrowheads) adjacent to the ventricular zone (VZ, brackets) rather than over the surface of the organoid. **b**. Laminin staining of enCORs following treatment with dissolved extracellular matrix (ECM). Note the presence of a laminin-rich basement membrane covering the surface of the organoid (arrow) and outside both the VZ (bracket) and newly generated neurons. **c**. Histological staining by haematoxylin and eosin reveals the presence of a radially oriented dense cortical plate (bracket) upon addition of dissolved ECM. **d**. Immunohistochemical staining for laminin and the neuronal markers MAP2 and Ctip2. Note the presence of remnant basement membrane without ECM addition (arrowheads), whereas upon addition of ECM a basement membrane (white arrows) forms outside the dense Ctip2+ cortical plate (yellow arrows). **e**. Nestin staining for radial glia reveals long basal processes (arrowheads) that terminate with end feet at the surface of the organoid (arrows) where the laminin-rich basement membrane (red) resides. **f**. Nuclear staining by DAPI reveals radially organized units (dashed lines) while sparse staining for radial glial fibers with the marker of mitotic radial glia Phosphorylated vimentin reveals fibers that extend the width of the tissue (arrowheads) reminiscent of radial units.

As a scaffold for migration and orientation, migrating neurons use the basal process of radial glial cells^31^, which contacts the basement membrane covering the surface of the brain. Staining for Nestin, a cytoplasmic marker of radial glia, revealed the presence of radial glial basal processes with end feet that terminated on the outer surface (Figure 3e). In contrast, spherical organoids displayed disorganized radial glial processes with terminating end feet at various locations within the tissue (Supplemental Fig. 10c). In addition, staining for phospho-vimentin, a cytoplasmic marker of dividing radial glia revealed long basal processes extending the length of the cortical wall (Figure 3f). Notably, nuclear staining alone revealed linear units of radial glia and neurons aligned in a manner reminiscent of radial units^31^, a characteristic architecture not previously recapitulated *in vitro*.

To verify the utility of the enCOR 3D culture method for disease modelling, we modelled Fetal alcohol syndrome (FAS), a leading preventable cause of intellectual disability affecting approximately 0.5-2 in 1000 births in the United States ^32^. FAS is characterized by neurodevelopmental abnormalities including microcephaly, thin or absent corpus callosum, and neuronal migration defects such as polymicrogyria, heterotopia, and focal lissencephaly ^33,34^. For this, 150mM or 300mM Ethanol (see methods) were included in the medium for 2 weeks beginning at day 46, and 60 day organoids were analysed for the presence of the cortical plate and the position of neurons. Strikingly, organoids treated with the lower concentration of ethanol displayed smaller cortical regions and a complete lack of a cortical plate (Figure 4a). Furthermore, neurons were located in abnormal clumps at the surface of the organoid consistent with the formation of external heterotopias, previously described in models of FAS ^35^. Organoids treated with the higher dose of ethanol displayed a large number of neurons abnormally located within the ventricular zone, having failed to migrate basally outward. This effect was so strong that it was difficult to identify a polarized ventricular zone in these tissues.

**Figure 4.**
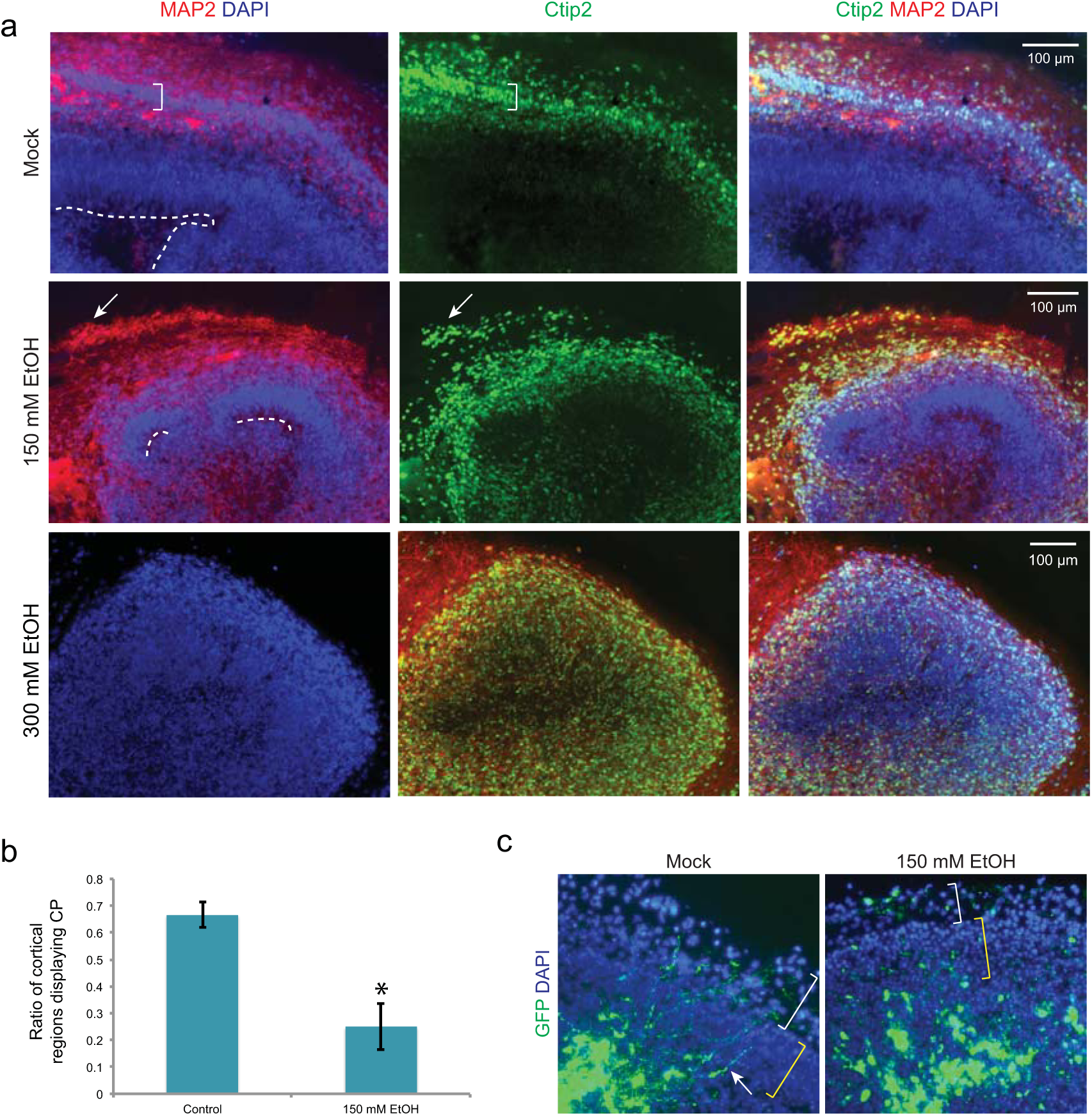
Ethanol treatment leads to neuronal migration defects and abnormal cortical plate formation. **a**. Treatment of enCORs with two doses of ethanol or water as mock control for two weeks reveals the loss of the dense cortical plate in ethanol treated organoids compared with control (bracket). Treated organoids display smaller ventricular surfaces (dashed lines) consistent with microcephaly, as well as overmigration of neurons (arrows). **b**. Quantification of the presence of a cortical plate reveals a dramatic reduction in number of cortical lobes containing a cortical plate in ethanol treated tissues. \**P*<0.05, Student’s *t*-test, n=5 organoids for control, 6 organoids for EtOH treatment. **c**. GFP labeled newborn neurons in electroporated tissues followed by ethanol treatment. Note the presence of long leading processes of neurons (arrow) migrating into the cortical plate (yellow brackets), whereas upon ethanol treatment neurons appear rounded and abnormal in morphology. White brackets denote the marginal zone.

Quantification of these findings revealed a significant reduction in cortical plate formation in organoids treated with 150 mM ethanol (Figure 4b). Furthermore, we performed electroporation of GFP in 67-day old organoids in order to mark newborn neurons and track their migration. On the day of electroporation, we began treatment with 150 mM ethanol and fixed and analysed the tissues 5 days later. Whereas control organoids displayed neurons with long leading processes migrating into the cortical plate, organoids treated with ethanol displayed abnormal neurons lacking these leading processes and failing to migrate into the cortical plate (Figure 4c). This effect of ethanol on the migrating neuronal leading process had not previously been described^36^ and could explain the disruption in formation of a radially organized cortical plate and development of heterotopias *in vitro* and *in vivo*.

The remarkable self-organization and the ability to generate the full repertoire of organ cell types have made organoids an important new model system. However, the high variability and difficulties modelling later tissue architecture has meant that subtle defects are difficult to discern. To overcome this, we have combined organoids with bioengineering using a novel microscale internal scaffold. This scaffold not only improves reproducibility, but also opens the door to future development such as testing of alternative materials, adsorption of instructive factors, and even gradient generation for axis formation. Finally, because enCORs show both a narrower range of variability and proper migration and formation of a cortical plate, the potentially discernable phenotypes are dramatically increased. This method enables the study of neuronal migration disorders such as lissencephaly, heterotopia, and polymicrogyria, as well as more subtle defects associated with autism, schizophrenia and epilepsy.

## Methods

### Preparation of microfilaments

Poly (lactide-co-glycolide) braided fibers of 10:90 PLGA were obtained commercially as Vicryl sutures (Ethicon). We preferentially used violet dyed fibers to assist in visualization during dispersion and within embryoid bodies. Individual microfilaments were isolated from the braided fiber by mechanical shearing with an angled blade against a stainless steel plate, to obtain filaments of 0.5-1mm in length, and 15μm in diameter. Filaments were then hydrated in embryoid body media and transferred to a 15ml conical tube for storage.

### Preparation of micropatterned embryoid bodies and enCORs

Embryoid bodies were prepared from single cell suspension of human ES or iPS cells, following accutase treatment, as described previously ^11^. Cells were counted and resuspended in embryoid body media (EB): DMEM/F12 (Invitrogen, cat. #11330-032) and 20% Knockout Serum replacement (Invitrogen, cat. #10828-028), 3% human ES quality batch-tested fetal bovine serum, 1% Glutamax (Invitrogen, cat. #35050-038), 1% MEM-NEAA (Sigma, cat. #M7145), 0.1 mM 2-mercaptoethanol, 4 ng/ml bFGF (Peprotech, cat. #100-18B), and 50uM Y-27632 ROCK inhibitor (VWR, cat. #688000-5). 18000 cells were added to each well of a 96-well low-attachment U-bottom plate (Sigma, cat. #CLS7007) already containing 5-10 microfilaments in embryoid body media, and media was added to give a final volume of 150ul per well. Based on an average hPSC cell size of 15 μm (measured from hPSC cell suspensions on EVOS microscope, Invitrogen), we calculated that 18000 cells per 5-10 fibers of maximum length 1mm would result in, at most, 5-10% of cells having direct contact to the fiber.

At day 3, half media was changed with EB media without bFGF and Y-27632. On day 5 or 6 depending on morphology, EBs were moved with an angled cut P200 tip to 24-well low-attachment plates (Sigma, cat. #CLS3473) with neural induction media (NI) as previously described ^11^. Media was changed every other day. On day 11, or when polarized neural ectoderm was visible on the surface, organoids were transferred to a droplet of Matrigel as previously described but kept in NI media. At day 13, media was changed to an improved differentiation media-A (IDM-A): 1:1 of DMEM/F12 and Neurobasal (Invitrogen, cat. #21103049), 0.5% N2 supplement (Invitrogen, cat. #17502048), 2% B27-vitamin A (Invitrogen, cat. #12587010), 0.25% insulin solution (Sigma, cat. #I9278-5ML), 50uM 2-mercaptoethanol, 1% Glutamax, 0.5% MEM-NEAA, and 1% Penicillin-Streptomycin (Sigma, cat. #P0781). Additionally, CHIR 99021 (Tocris, cat. #4423) at 3uM was added from day 13 to 16. Media was changed every other day and organoids were moved to a spinning bioreactor or orbital shaker as described previously, on day 18.

Spherical organoids were generated exactly as described previously ^11^. For testing the effects of early growth factors on EB neural induction, 3N media containing retinoids and dual-SMAD inhibitors was prepared as previously described ^15^.

### Formation of polarized cortical plate

At day 20, media was changed to an improved differentiation +A (IDM+A): 1:1 of DMEM/F12 and Neurobasal, 0.5% N2 supplement, 2% B27 +vitamin A, 0.25% insulin solution, 50uM 2-mercaptoethanol, 1% Glutamax, 0.5% MEM-NEAA, 1% Penicillin-Streptomycin, 0.4 mM Vitamin C, and 1.49g HEPES per 500 ml to control pH levels. Alternatively, media can be pH controlled with further bicarbonate buffering with the addition of 1mg/ml sodium bicarbonate. At day 40, media was changed to IDM+A with 1ml dissolved Matrigel per 50 ml media by slowly thawing the Matrigel on ice and addition to cold media to dissolve. For ethanol treatments, ethanol was added at a final concentration of 150mM or 300 mM or an equal volume of water was used as control mock treated. These concentrations are higher than physiological levels but given the volatility of ethanol, this has been shown to be the most effective in eliciting an effect in neurons *in vitro* and in brain slice culture^37,38^. The media was changed with new media containing fresh ethanol every 3-4 days.

### Histological and immunohistochemical analysis

Organoids were fixed in 4% paraformaldehyde for 20 min at room temperature and washed with PBS three times for 10 min each at room temperature before allowing to sink in 30% sucrose at 4°C. The tissues were embedded and sectioned and stained as described^11^. Primary antibodies used were: Brachyury (R and D Systems AF2085, 1:200), mouse anti-N-Cadherin (BD 610920, 1:500), mouse anti-E-Cadherin (BD 610182, 1:200), goat anti-Sox17 (R and D systems AF1924, 1:200), rabbit anti-Laminin (Sigma L9393, 1:500), rabbit anti-Tbr1 (Abcam ab31940, 1:300), chicken anti-Tbr2 (Millipore AB15894, 1:100), mouse anti-Map2 (Chemicon MAB3418, 1:300), rat anti-Ctip2 (Abcam, ab18465, 1:300), rabbit anti-Arl13b (Proteintech 17711-1-AP, 1:300), mouse anti-phospho-Vimentin (MBL International D076-3S, 1:250), rabbit anti-Emx1 (Sigma HPA006421, 1:200). DAPI was added to secondary antibody to mark nuclei. Secondary antibodies labeled with Alexafluor 488, 568, or 647 (Invitrogen) were used for detection. For histological analysis, sections were stained for hematoxylin/eosin followed by dehydration in ethanol and xylene and mounting in permount mountant media. Images were acquired on a confocal microscope (Zeiss LSM 710 or 780). Statistical analysis of imaging data was performed using Student’s *t*-test for significance.

### Electroporation of organoids

Electroporation of pmax-GFP construct (Lonza) was performed as described previously^19^ and organoids were immediately treated with 150 mM ethanol or an equal volume of water as control for 5 days before PFA fixation and embedding and sectioning as described above. Sections were stained for DAPI and imaged immediately.

### RT-PCR analysis of gene expression

Three organoids for each condition were collected in Trizol reagent (Thermo Fisher) and RNA isolated according to manufacturer. DNA was removed using DNA-Free kit (Ambion) and reverse strand cDNA synthesis was performed using Superscript III (Invitrogen). PCR was performed using primers for a panel of pluripotent and germ layer identities (R and D systems, SC012).

### RNA-seq analysis

Three individual organoids for each condition were collected at the indicated time points. RNA was isolated using Arcturus PicoPure RNA Isolation Kit (Thermo Fisher Scientific, cat. #KIT0204) (20 days timepoint) or Trizol Reagent (Thermo Fisher Scientific, cat. # 15596018) (60 days timepoint) according to the manufacturers instructions. RNA concentration and integrity was analysed using RNA 6000 Nano Chip (Agilent Technologies, cat. #5067-1511). RNA was enriched for mRNA using Dynabeads mRNA Purification Kit (Thermo Fisher Scientific, cat. # 61006). Libraries were prepared using NEBNext Ultra Directional RNA Library Prep Kit for Illumina (NEB, cat. # E7420L). Barcoded samples were multiplexed and sequenced 50bp SE on a HighSeq2500 (Illumina). Sample preparation and sequencing was performed at the VBCF NGS Unit (www.vbcf.ac.at). Data will be submitted to GEO and accession numbers provided.

The strand specific reads were screened for ribosomal RNA by aligning with BWA (v0.6.1)^39^ against known rRNA sequences (RefSeq). The rRNA subtracted reads were aligned with TopHat (v1.4.1)^40^ against the Homo sapiens genome (hg38) and a maxiumum of 6 missmatches. Maximum multihits was set to 1 and InDels as well as Microexon-search was enabled. Additionally, a gene model was provided as GTF (UCSC, 2015_01, hg38). rRNA loci are masked on the genome for downstream analysis. Aligned reads are subjected to FPKM estimation with Cufflinks (v1.3.0)^41,42^. In this step bias detection and correction was performed. Furthermore, only those fragments compatible with UCSC RefSeq annotation (hg38) of genes with at least one protein-coding transcript were allowed and counted towards the number of mapped hits used in the FPKM denominator. Furthermore, the aligned reads were counted with HTSeq (0.6.1p1) and the genes were subjected to differential expression analysis with DESeq2 (v1.6.3)^43^.

GO term enrichment analysis was performed on genes with an adjusted p value <0.1 and absolute log2fc value >1 in at least one of the conditions (20 day or 60 day, spheroid or enCOR). Each set of differentially expressed genes were analysed using Pantherdb.org for GO Slim biological process enrichment and fold enrichment for terms was plotted (Supplementary Table 1).

For hierarchical cluster analysis, genes were reorganized based on their similarity of log2fc values by means of Ward hierarchical clustering using heatmap.2 of the gplots package in R. Cutree function gave the 7 clusters used for subsequent GO term analysis. Gene lists were fed into Pantherdb.org for GO Biological Process Complete, yielding a large list of redundant terms. Therefore, in order to remove redundancy, we narrowed the list using GO Trimming and a cutoff of terms with fold enrichment value >2 yielding the list of GO terms that were plotted by fold enrichment value (Supplementary Table 2). Individual tracks were visualized using Interactive Genomics Viewer IGV_2.3.68 (Broad Institute).

For comparison to Allen BrainSpan human transcriptome dataset, the RPKM expression values of Brainspan were downloaded (http://www.brainspan.org/static/download.html). FPKM values were filtered for differential expression in the 60d time-points (padj < 0.1) and joined with Brainspan via the gene symbols. The similarity of expression was compared by the rank of the expressed gene via Spearman correlation. The heatmap of Spearman coefficient for each region at fetal time points was generated using heatmap.2 without hierarchical clustering. The list was manually ordered according to anterior-posterior regional position and separated into four developmental stages.

## Acknowledgements

We thank members of the Knoblich lab for insight and technical help, particularly Simone Wolfinger, Hilary Gustafson and Angela Peer. M.A.L. was funded by a Marie Curie Postdoctoral fellowship, N.C. was funded by EMBO and DFG postdoctoral fellowships. Work in M.A.L.’s laboratory is supported by the Medical Research Council MC_UP_1201/9. Work in J.A.K.’s laboratory is supported by the Austrian Academy of Sciences, the Austrian Science Fund (grants I_1281-B19 and Z_153_B09), and an advanced grant from the European Research Council (ERC).

## Author contributions

M.A.L. conceived the project, planned and performed experiments and wrote the manuscript. N.S.C. performed RNA library preparation and analyzed data. T.R.B. performed bioinformatics analysis of RNAseq data. J.A.K. supervised the project, planned and interpreted experiments, and wrote the manuscript.

---

<u>Supplemental Figure 1. Batch-to-batch variability in neuroectoderm formation in spherical organoids</u> Bright field images of five independent batches of spheroid organoids showing the degree of variability in generation of polarized neuroectoderm and quantification at the right.

<u>Supplemental Figure 2. Non-neural tissue formation in spherical organoids</u> **a**. Immunohistochemical staining of day 10 spheroids for the germ layer markers Brachyury for mesoderm, N-Cadherin for neural ectoderm, Sox17 for endoderm and E-Cadherin for non-neural epithelium. Note the polarized neural epithelia (arrows) displaying the apical domain on the surface (white asterix) on an optimal brain organoids, while extensive mesoderm and endoderm identities (arrowheads) are visible in suboptimal organoids. **b**. Haematoxylin and eosin staining of spheroid organoids showing lobes of brain tissue (arrows) but also non-neural regions such as fluid-filled cysts and fibrous regions (arrowheads). **c**. Immunohistochemical staining of a spheroid organoid reveals occasional endoderm (Sox17+, inset) and nonneural epithelia (arrow) even at a later time point of 40 days.

<u>Supplemental Figure 3. Dual-SMAD+retinoid treatment of spherical organoids compromises tissue integrity</u> **a**. Bright field images showing that treatment of spherical EBs with dual-SMAD inhibition and retinoids improves early (day 3) morphology with a greater extent of surface clearing. **b**. Bright field imaging following matrigel embedding, however, reveals that treated EBs do not form large buds of neural epithelium (arrows in control) instead generating numerous small rosettes and already generating neurons with neurites (arrowheads).

<u>Supplemental Figure 4. Micropatterned brain organoids display consistent generation of neural ectoderm</u> **a**. Bright field image of the intact braided PLGA fiber. **b**. Bright field image of isolated microfilaments. **c**. Hydrated microfilaments within a droplet of EB medium. **d**. Three examples of micropatterned EBs at day 3 of the protocol. Note the elongated and sometimes complex arrangement. **e**. Bright field images of five independent batches of microfilament patterned organoids showing reduced variability in generation of polarized neuroectoderm and quantification at the right.

<u>Supplemental Figure 5. Consistent generation of pure neural identity in enCORs</u> **a**. Immunohistochemical staining for Sox17 for endoderm and E-Cadherin for non-neural epithelium reveals only occasional cells (arrowhead) with these non-neural identities. The microfilament can be seen as an autofluorescent rod (yellow asterisk). **b**. RT-PCR for markers of pluripotency in microfilament patterned organoids or spherical organoids. Embryonic stem cells are positive control and Neg. is the water control. **c**. RT-PCR for expression of markers of the three germ layers: Neuroectoderm (NE), mesoderm (ME), and endoderm (EN) in microfilament patterned organoids and organoids lacking a filament (spherical organoids). Neg. is the negative water control. **d**. Bright field images several days following matrigel embedding showing the prevalence of non-neural cysts in unpatterned organoids compared with microfilament patterned organoids, and quantification at the right. \**P*<0.05, Student’s *t*-test, n=97 organoids from 5 batches for spheroid, n=69 organoids from 5 batches for micropatterned.

<u>Supplemental Figure 6. Reproducible formation of dorsal forebrain tissue in enCORs with pulse of CHIR</u> **a**. Bright field image of micropatterned organoids shortly after matrigel embedding, displaying numerous small buds of neuroepithelium (arrows). **b**. Embedded micropatterned organoids following three days of treatment with CHIR. Note the larger, more continuous buds of neuroepithelium. **c**. Histological section stained with haematoxylin and eosin showing pure neural tissue and many large continuous lobes of brain tissue (arrows). **d**. Immunohistochemical staining for Sox17 and E-Cadherin showing an absence of nonneural identities and several large lobes of brain tissue (arrows) in micropatterned organoids. **e**. Immunohistochemical staining for the dorsal cortical marker Emx1 (green) and the neural identity marker N-Cadherin (red) reveals much of the tissue is composed of dorsal cortical brain regions (arrows). **f**. Staining of micropatterned +CHIR brain organoids and spheroid organoids for the markers of dorsal cortex Tbr1 and Tbr2 reveals all large lobes of tissue are dorsal cortex (arrows) in micropatterned organoids. Organoids that were not patterned (spheroid) show much fewer dorsal regions and large brain regions that lack this identity.

<u>Supplemental Figure 7. Generation of a predominantly dorsal forebrain tissue in enCORs.</u> **a**. Staining for the marker Foxg1 reveals the presence of large forebrain regions in both patterned and spheroid organoids, but a lack of the marker Pax6 in this region of a spheroid. **b**. Staining for the ventral forebrain marker Gsh2 and the dorsal marker Tbr2 reveals the presence of large ventral regions in the unpatterned organoids whereas micropatterned +CHIR organoid display mainly large Tbr2+ dorsal regions with occasional smaller Gsh2 ventral regions. **c**. Staining for the marker of choroid plexus TTR reveals the presence of this region in both patterned and unpatterned organoids. **d**. Staining for the marker of hippocampus Prox1 reveals the presence of this tissue in both spheroid and micropatterned organoids. **e**. Staining for the spinal cord marker Pax2 reveals staining in regions of a spheroid organoid, but a lack of staining in the micropatterned organoid.

<u>Supplemental Figure 8. GO terms in identified clusters of gene expression</u> Fold enrichment of GO Biological Process terms for genes in 7 clusters identified by hierarchical clustering according to log2fc value (Figure 2e).

<u>Supplemental Figure 9. Individual marker genes of pluripotency, germ layer identity and brain patterning</u> **a**. Single gene tracks of markers of pluripotency, neuroectoderm and mesendoderm in spheroid and micropatterned organoids at 20 days. **b**. Single gene tracks of markers of rostral-caudal and dorsal-ventral brain patterning in spheroid and micropatterned organoids at 60 days. Schematic of marker expression in the developing brain shown below.

<u>Supplemental Figure 10. enCORs with ECM addition display organized cortical plate while spheroids display a disorganized radial scaffold</u> **a**. Bright field images of organoids lacking dissolved ECM and microfilament patterned organoids with dissolved ECM showing a band of density upon addition of ECM (arrow) reminiscent of a dense cortical plate. **b**. Higher magnification of immunostaining for Laminin, MAP2 and Ctip2 showing the radial orientation of neurons that have reached the cortical plate (brackets). **c**. Nestin staining in spheroid organoids reveals the presence of radial glial processes that show disorganization outside the ventricular zone (arrowheads) and terminal end feet within the tissue (arrows).

<u>Supplementary Table 1</u> Upregulated and downregulated genes in micropatterned organoids at 20 days and 60 days, and the associated GO term analysis.

<u>Supplementary Table 2</u> Hierarchical clusters of up and down regulated genes and the associated GO term analysis.

